## Supplemental figures for "The critical role of enterovirus 2A protease in viral translation, replication, and antagonism of host antiviral responses"

### **Supplementary figures**

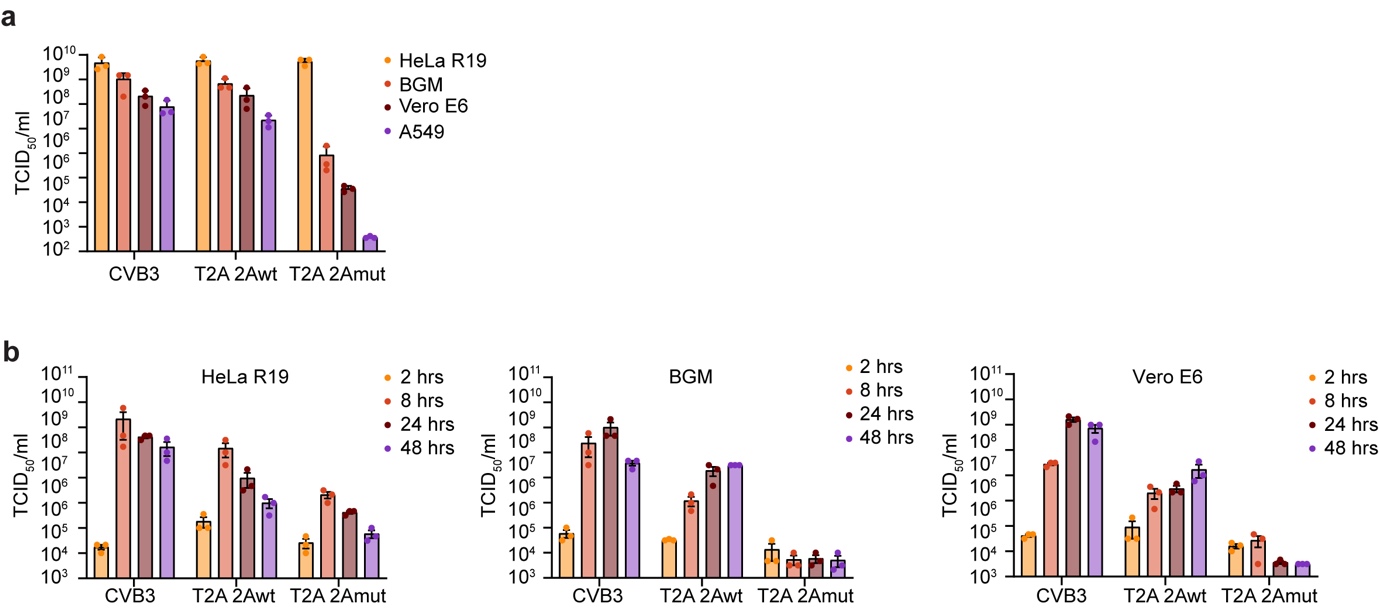

**Supplementary Fig. 1: Growth kinetics of 2Awt and 2Amut viruses on different cell lines.** (**a**) CVB3, 3CD_cs_-2Awt, and 3CD_cs_-2Amut viral stocks were titrated on HeLa-R19, BGM, Vero E6 and A549 by end-point dilution. (**b**) Growth curves of CVB3, T2A-2Awt and T2A-2Amut viruses in HeLa-R19, BGM, Vero E6 and A549 cells performed as in Fig.1e. Data represent the mean ± SEM of the three technical replicates (**a**,**b**).

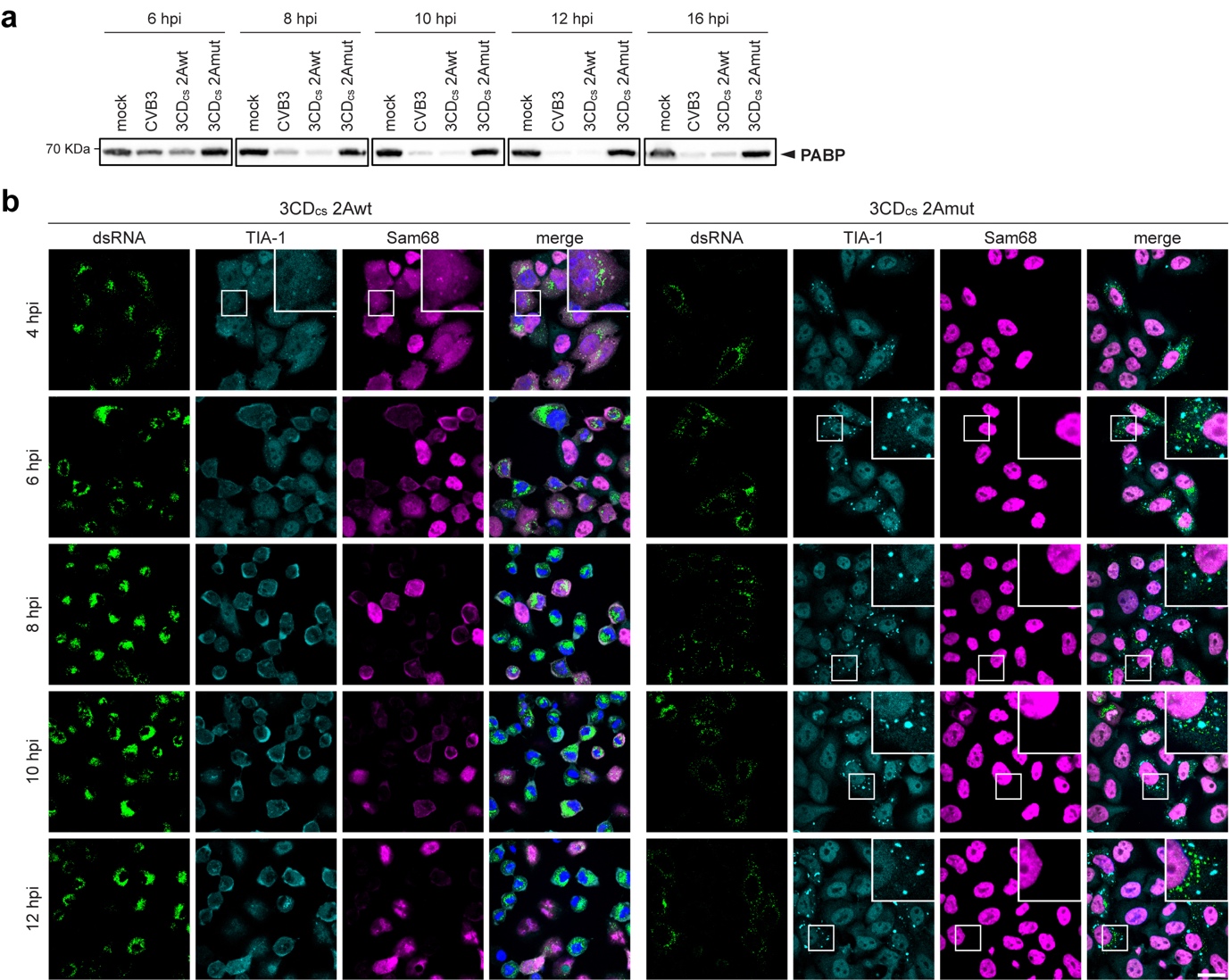

**Supplementary Fig. 2: Cleavage of PABP at later time points and SGs in infected cells stained with TIA-1 and Sam68.** (**a**) Western blot analysis for PABP. Cell lysates derive from the same experiment shown in Fig. 4d. (**b**) IF analysis of HeLa-R19 cells infected with CVB3, 3CD_cs_-2Awt and 3CD_cs_-2Amut viruses, fixed and stained for dsRNA as infection marker and TIA-1 and Sam68 as SG marker. Experiment was performed as described in Fig. 4c. Magnified regions are shown in white boxes. Scale bar: 25 μm.

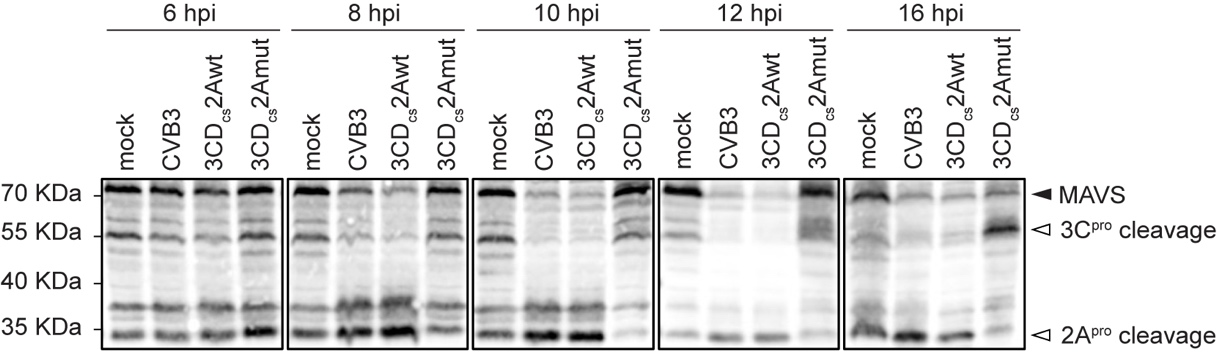

**Supplementary Fig. 3: MAVS cleavage products generated by 3C^pro^ and 2A^pro^ in CVB3-2Awt and CVB3-2Amut infected cells.** Western-blot analysis of MAVS in cell lysates obtained from the same experiment shown in Fig. 4d. Black triangles denote intact proteins, white triangles the cleavage products.

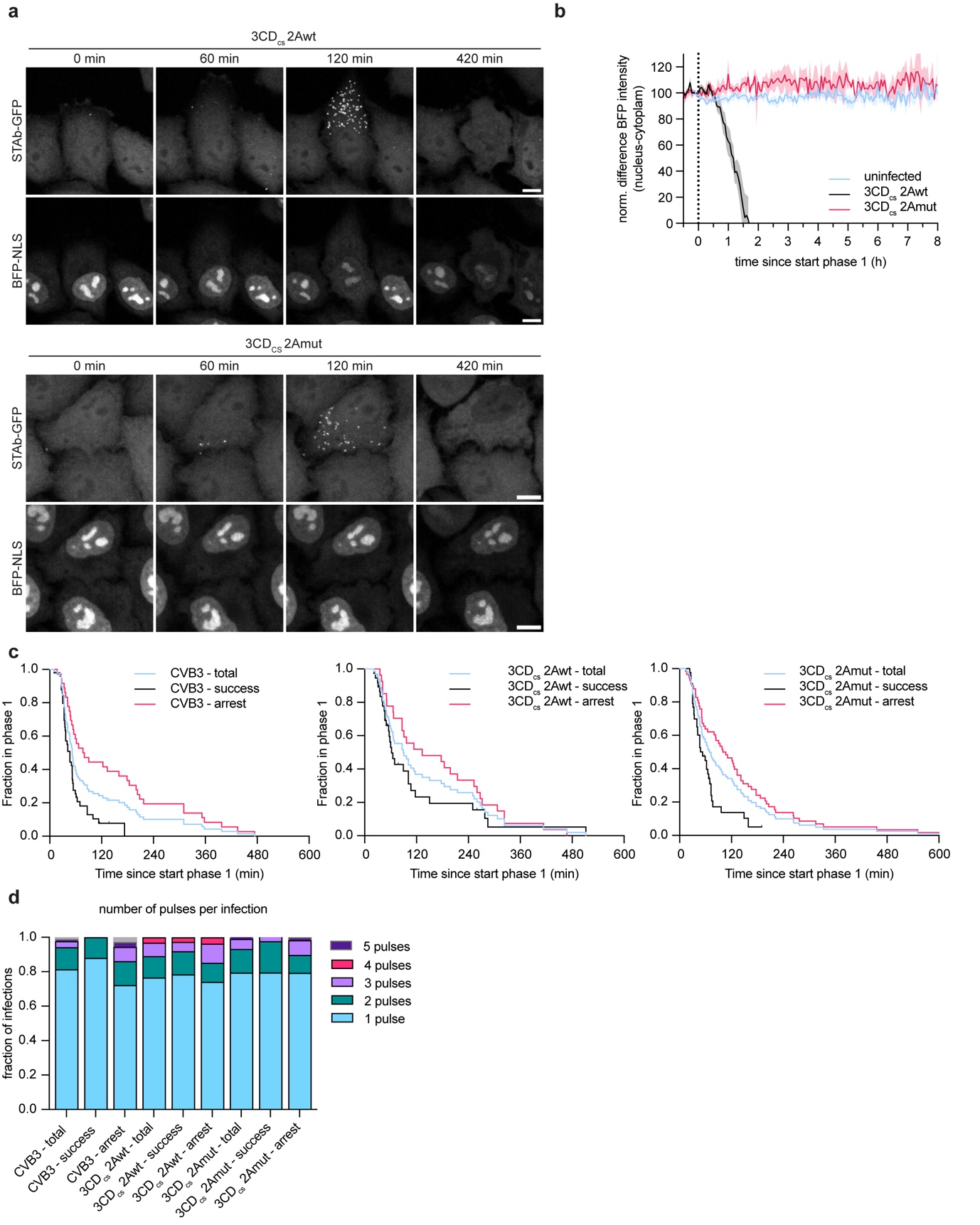

**Supplementary Fig. 4: 2A^pro^ rapidly and exclusively triggers nucleocytoplasmic trafficking disorder.** Hela-R19 saGFP-STAb BFP-NLS C1 cells were infected at an MOI of 0.25 with ST-CVB3-2Awt or ST-CVB3-2Amut and time-lapse imaging of STAb-GFP and BFP-NLS was performed. (**a**) Representative pictures of cells infected with ST-CVB3-2Awt or -2Amut at different time points aligned to the start of phase 1. (**b**) average normalized BFP-NLS intensity ratio between nucleus and cytoplasm over time aligned to start phase 1. Scale bar: 10 μm. (**c**) Kaplan-Meier curves depicting duration of phase 1 in those cells with successful replication, non-successful replication (i.e., phase 2 arrested), or both (total). (**d**) Fraction of infections with indicated number of translation pulses.

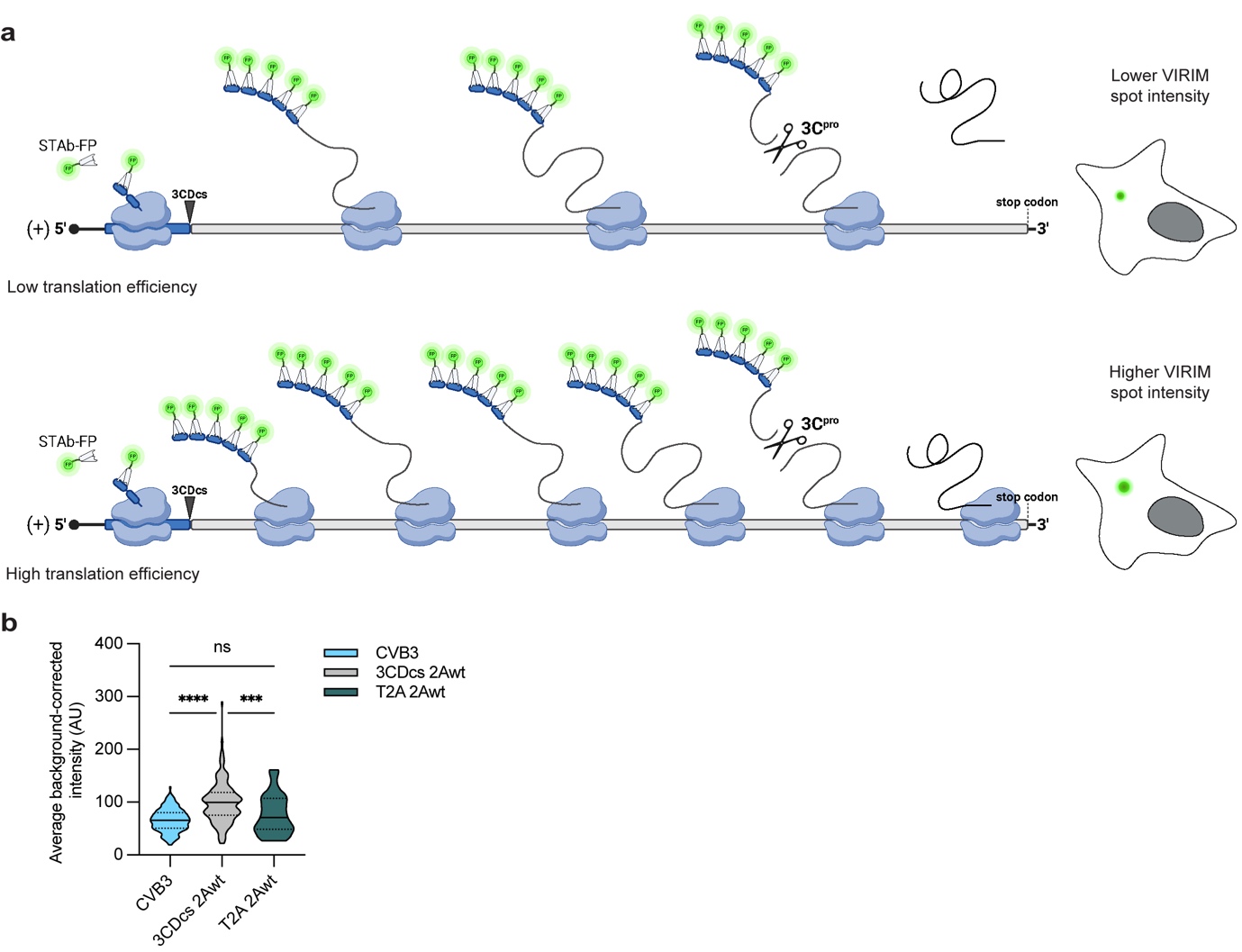

**Supplementary Fig. 5: VIRIM spot intensity as a measure of viral translation efficiency.** (**a**) Schematic representation of VIRIM spot intensity analysis. Upper panel shows a situation with comparatively lower translation efficiency, less translating ribosomes on a single vRNA, and consequently a smaller accumulation of fluorescence signal. The lower panel depicts the scenario of a comparatively higher translation efficiency. (**b**) Absolute intensity measurements of different CVB3 mutants. The higher intensity for 3CDcs-2Awt is the consequence of a relatively longer effective transcript length (i.e., the length of transcript that is decoded before the nascent peptide chain is released from the ribosome). statistical significance was assessed by a one-way analysis of variance (ANOVA) with multiple comparisons testing. * p<0.5 ; ** p<0.01 ; *** p<0.001.
